## Supplementary material for "*Rickettsia* infection rate along an altitudinal gradient as influenced by population genetic structure of Ixodid ticks": S2 Table

**Supplementary Table 1.** The difference in *Rickettsia* infection rate in *H. flava* haplotype groups. Table of the distribution of *Rickettsia* infected and uninfected *H. flava* in each of the genetic groups as shown in Supplementary Figure 2 *(H. flava* median joining network)*.* The results of the z-score test for two populations proportions at p<0.05 showed no significant difference between the *Rickettsia* detection rate in haplotype groups 1 and 2. Welch t-test at p<0.05 revealed no significant difference in the mean altitude of the two groups.

| Haplotype group | *Rickettsia* infection | | | Mean altitude |
| --- | --- | --- | --- | --- |
|  | Positive | Negative | Detection rate |  |
| 1 | 1 | 45 | 2.17%^ab^ | 205^ab^ |
| 2 | 14 | 183 | 7.11%^ab^ | 165^ab^ |
