## Supplementary material for "*Rickettsia* infection rate along an altitudinal gradient as influenced by population genetic structure of Ixodid ticks": S3 Figure

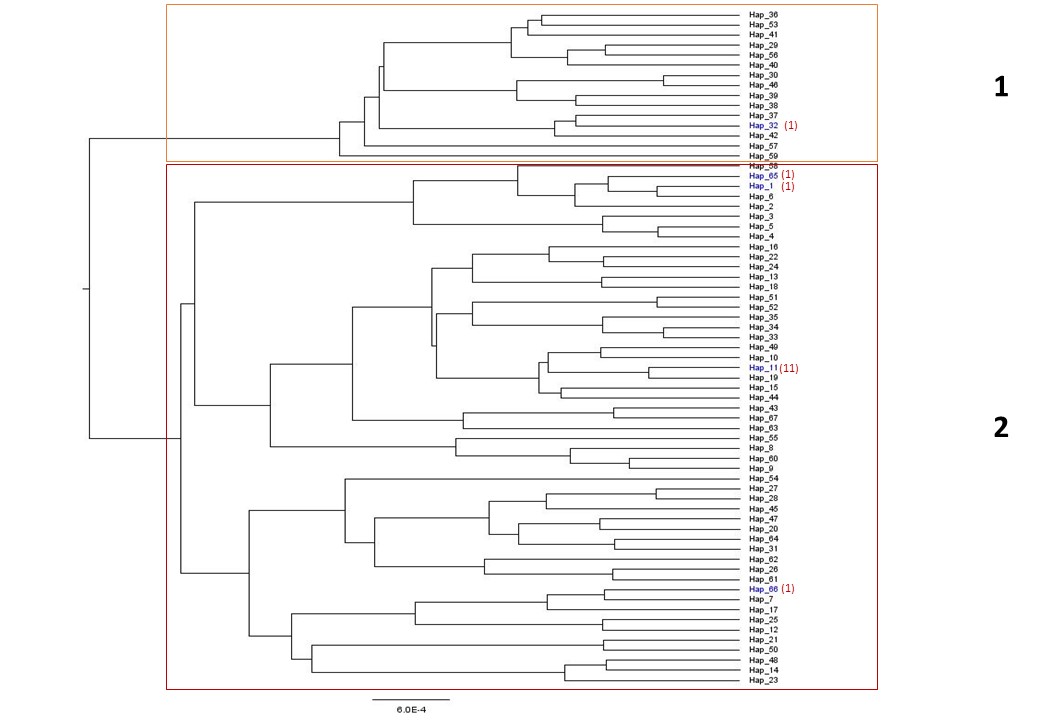

**Supplementary Figure 3.** Phylogenetic tree from BEAST analysis of 66 haplotype *cox1* sequences of *H. flava.* The blue labelled haplotypes indicate *Rickettsia* infection in individual samples. The parenthesis in red shows the number of *Rickettsia* infected tick. The black labelled haplotypes are negative for *Rickettsia* infection.
