## Supplementary material for "*Rickettsia* infection rate along an altitudinal gradient as influenced by population genetic structure of Ixodid ticks": S1 Figure

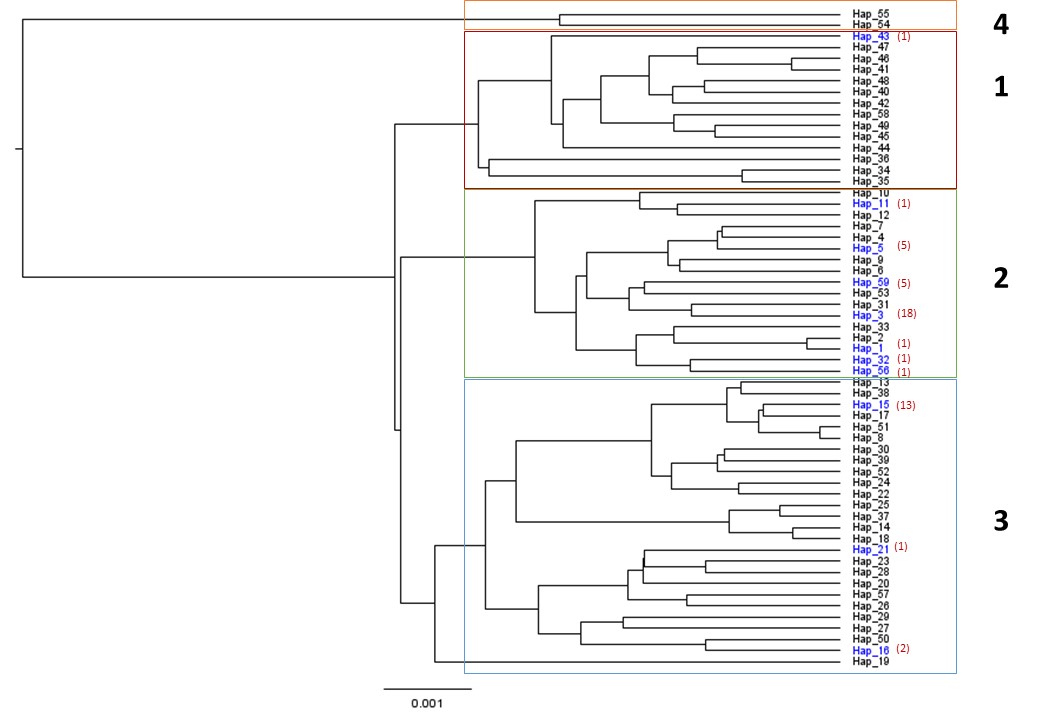

**Supplementary Figure 1.** Phylogenetic tree from BEAST analysis of 59 haplotype *cox1* sequences of *I. ovatus.* The blue labelled haplotypes indicate the presence of *Rickettsia* infection. The red parenthesis beside is the number of *Rickettsia* positive individuals per haplotype. The black labelled haplotypes are negative for *Rickettsia* infection.
