## Supplementary material for "*Rickettsia* infection rate along an altitudinal gradient as influenced by population genetic structure of Ixodid ticks": S2 Figure

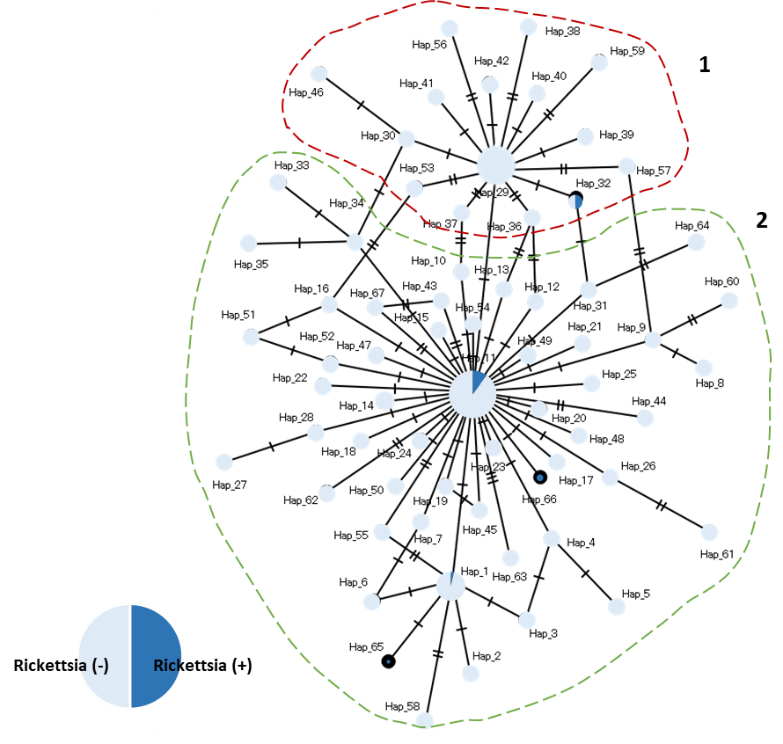

**Supplementary Figure 2.** Median joining network of the 66 *cox1* haplotype sequences of *Rickettsia* positive and negative *H. flava.* Haplotype groups are indicated as 1 and 2.
